# The role of mesotocin on social bonding in pinyon jays

**DOI:** 10.1101/599555

**Authors:** Juan F. Duque, Tanner Rasmussen, Anna Rodriguez, Jeffrey R. Stevens

## Abstract

The neuropeptide oxytocin influences mammalian social bonding by facilitating the building and maintenance of parental, sexual, and same-sex social relationships. However, we do not know whether the function of the avian homologue mesotocin is evolutionarily conserved across birds. While it does influence avian prosocial behavior, mesotocin’s role in avian social bonding remains unclear. Here, we investigated whether mesotocin regulates the formation and maintenance of same-sex social bonding in pinyon jays (*Gymnorhinus cyanocephalus*), a member of the crow family. We formed squads of four individually housed birds. In the first, ‘pair-formation’ phase of the experiment, we repeatedly placed pairs of birds from within the squad together in a cage for short periods of time. Prior to entering the cage, we intranasally administered one of three hormone solutions to both members of the pair: mesotocin, oxytocin antagonist, or saline. Pairs received repeated sessions with administration of the same hormone. In the second, ‘pair-maintenance’ phase of the experiment, all four members of the squad were placed together in a large cage, and no hormones were administered. For both phases, we measured the physical proximity between pairs as our proxy for social bonding. We found that, compared to saline, administering mesotocin or oxytocin antagonist did not result in different proximities in either the pair-formation or pair-maintenance phase of the experiment. Therefore, at the dosages and time frames used here, exogenously introduced mesotocin did not influence same-sex social bond formation or maintenance. Like oxytocin in mammals, mesotocin regulates avian prosocial behavior; however, unlike oxytocin, we do not have evidence that mesotocin regulates social bonds in birds.

## Introduction

A group of young male pinyon jays fly from pine tree to pine tree consuming seeds as they go. Two of the birds are inseparable, never straying more than a few feet from each other. Other jays come and go from the group, but this dyad stays together for the season, even though they are not related. This dyad shares a strong bond, and each member of the dyad has weaker bonds with other individuals. Similar patterns occur in the interactions when humans engage in social events. Although everyone is together, the sociality of individuals varies. Some congregate in tight groups “catching up”, while others remain separate from groups, sticking near the food bar or off to the side of the room.

Having strong social connections is beneficial to survival and reproduction (Silk, 2007; Clutton-Brock, 2016). For example, maternal behavior depends on the bond created after birth and during nursing in mammals, particularly in species that give birth to a single offspring at a time rather than a litter (Broad, Curley, & Keverne, 2006). Notably, the maternal behaviors—nursing, grooming, and infant retrieval—are essential to the health and survival of the offspring and thus reproductive success of the mother. Further, strong female-female bonds often lead to maternal behavior by females other than the offspring’s mother, which are critical to the survival and reproduction of the offspring (Hrdy, 1999; Broad, Curley, & Keverne, 2006). Long-term study of savannah baboons has shown sociality and individual bonds between females to lead longer female longevity and increased infant survival (Silk, Alberts, & Altmann, 2003; Silk, Beehner, Bergman, Crock-ford, Engh, Moscovice, Wittig, Seyfarth, & Cheney, 2010). In feral horses, these female-female bonds benefit both the survival of individual foals and overall fecundity of the mares involved. In fact, these bonds seem to limit harmful behavior in the males, such as aggression toward mares, harassment, and infanticide (Cameron, Setsaas, & Linklater, 2009).

Social bonds provide obvious adaptive benefits, but what physiological mechanisms underlie these bonds? The neuropeptide hormone oxytocin (OT) plays a key role in a range of social behaviors. For example, sharing food increases levels of oxytocin circulating in the body of chimpanzees (Wittig, Crockford, Deschner, Langergraber, Ziegler, & Zuberbühler, 2014), and administering oxytocin to dogs increases gazing behavior at owners (Nagasawa, Mitsui, En, Ohtani, Ohta, Sakuma, Onaka, Mogi, & Kikusui, 2015). Further, oxytocin regulates the development of pair bonds and mother-offspring bonds. In rats, maternal behaviors, such as nursing and infant retrieval, act as a positive feedback for both mother and pups, resulting in increasing levels of oxytocin that strengthen their attachment (Nagasawa, Okabe, Mogi, & Kikusui, 2012). Administering oxytocin can induce similar maternal behavior in sheep that do not have offspring (Costa, Guevara-Guzman, Ohkura, Goode, & Kendrick, 1996). In the prairie vole, a primarily monogamous species, administration of oxytocin to females can establish mating pair and maternal bonds, whereas administration of an oxytocin antagonist can hinder such bonds (Insel, Winslow, Wang, & Young, 1998). In female marmosets, oxytocin administration induces greater preference for the male they were previously paired with and seems to make individuals in established bonded-pairings less likely to form social bonds with opposite sex strangers (Cavanaugh, Mustoe, Taylor, & French, 2014).

Oxytocin also plays a key role in mammalian social bonds among unrelated individuals outside of the pair bond. However, it remains unclear how oxytocin regulates these bonds. In humans, oxytocin levels can affect trust between non-kin humans (Kosfeld, Heinrichs, Zak, Fischbacher, & Fehr, 2005; Baumgartner, Heinrichs, Vonlanthen, Fischbacher, & Fehr, 2008), though its effects depend on context (Bartz, Zaki, Bolger, & Ochsner, 2011; Nave, Camerer, & McCullough, 2015). In chimpanzees, oxytocin levels increase when socially bonded partners groom but not when non-bonded partners groom (Crockford, Wittig, Langergraber, Ziegler, Zuberbühler, & Deschner, 2013). Oxytocin plays a complicated role in capuchin monkey social proximity, with oxytocin administration actually increasing social distance rather than decreasing it (Brosnan, Talbot, Essler, Leverett, Flemming, Dougall, Heyler, & Zak, 2015; Benítez, Sosnowski, Tomeo, & Brosnan, 2018). Despite these data, we do not understand how oxytocin underlies the initial formation of the mammalian social bond itself and, then, once a bond is established, the role that it plays in maintaining that social bond.

Here, we sought to assess the role of oxytocin in social bond formation and maintenance. We investigated this in pinyon jays (*Gymnorhinus cyanocephalus*), a highly social North American corvid. Like many social primates, pinyon jays have a fission-fusion-like dynamic social system in which individuals are typically part of a small, tight-knit sub-group of 5-20 individuals, but sub-groups often congregate, forming large flocks of up to 500 individuals (Marzluff & Balda, 1992). Juveniles commonly engage in social play and associate with many others, and as they become yearlings and early adults begin to preferentially associate with sub-group members and their pair-bonded partners (Marzluff & Balda, 1992). However, new associations continue throughout life as pinyon jays exhibit a fission-fusion-like social structure and are thus exposed to many new individuals, including opportunities for cooperation. For example, unrelated adult pinyon jays engage in prosocial behavior, particularly through the sharing of food. Though food sharing between same-sex pairs of birds is not dependent on reciprocity, more dominant birds may be more likely to share with subordinate ones, which suggests sharers may be receiving social benefits (Duque & Stevens, 2016). Moreover, administering mesotocin (MT), the avian homologue to oxytocin, increases the likelihood that pinyon jays will voluntarily be generous to others. If given an option between providing food for only itself or itself and another individual (prosocial choice), mesotocin increases the preference for the prosocial action (Duque, Leichner, Ahmann, & Stevens, 2018). Thus, the long-lived and highly social nature of pinyon jays and evidence of mesotocin influencing their prosociality make them ideal candidates to study how social bonds form.

Both oxytocin and mesotocin are nine amino acid peptides but mesotocin has a minor amino acid substitution from leucine to iso-leucine in position 8 (Acher, Chauvet, & Chauvet, 1970). Mesotocin seems to be a functional homologue to oxytocin in birds because its administration increases preferences for larger over smaller social groups (Goodson, Schrock, Klatt, Kabelik, & Kingsbury, 2009) and increases prosocial preferences (Duque, Leichner, Ahmann, & Stevens, 2018), whereas administering an antagonist disrupts pair bond formation (Pedersen & Tomaszycki, 2012). Therefore, we aimed to assess mesotocin’s role in social bond formation and maintenance in birds.

Our first research question investigated whether mesotocin is critical to the formation of social bonds among unrelated, same-sex pinyon jays. We tested this by administering mesotocin, an oxytocin antagonist, or saline to previously unfamiliar pairs of individuals in repeated interactions. The short-term effects of this hormone on social bonds were assessed by measuring the proximity between individuals and comparing these distances across hormone conditions. If mesotocin builds social bonds, repeated exposure to mesotocin when paired with a particular individual should create a strong bond as measured by proximity. Exposure to oxytocin antagonist or saline should produce weaker or no bonds.

Our second research question investigated whether mesotocin provides long-term social bond maintenance in a group. We tested this by placing the pairs in larger groups in the absence of further hormone administration and measuring proximity between all group members. If mesotocin enhances the initial formation of a relationship between two individuals, then those bonds should remain when multiple individuals are present in a group, even without further mesotocin administration. Conversely, pairs treated with either oxytocin antagonist or saline should show less social proximity in the group setting.

## Methods

### Subjects

We conducted two experiments with independent sets of adult pinyon jays: 12 birds (8 male, 4 female) in Experiment 1 from September to December 2015 and 24 birds (16 male, 8 female) in Experiment 2 from September to December 2017. Researchers captured all birds as wild adults in either Arizona or California (USFW permit MB694205) between 1996 and 2011. All birds were housed in individual cages since capture and placed in one of three housing rooms where they had visual and acoustic contact with other jays in their room. Housing rooms were kept at 22° C with a 14:10 h light:dark cycle. Birds were fed Lafeber’s Cockatiel and Parrot Pellets, turkey starter, live mealworms, pine nuts, and peanuts daily. The University of Nebraska-Lincoln IACUC approved this project (protocols 834 and 1354) and all procedures conformed to the ASAB/ABS Guidelines for the Use of Animals in Research.

### Formation of squads and pairs

We assigned each subject to a same-sex squad of four individuals (three squads in Experiment 1 and six squads in Experiment 2). When possible, squads were formed from individuals within the same housing room (47 of all 54 pairs were from the same housing room). Thus, most pairs should have been familiar with each other. Within these squads, we paired subjects with each of the three other individuals in the same squad, creating six total pairs per squad (Figure 1). All birds were individually housed when not being run through experimental sessions, so birds only had direct experience with squad members during experimental sessions.

**Figure 1.**
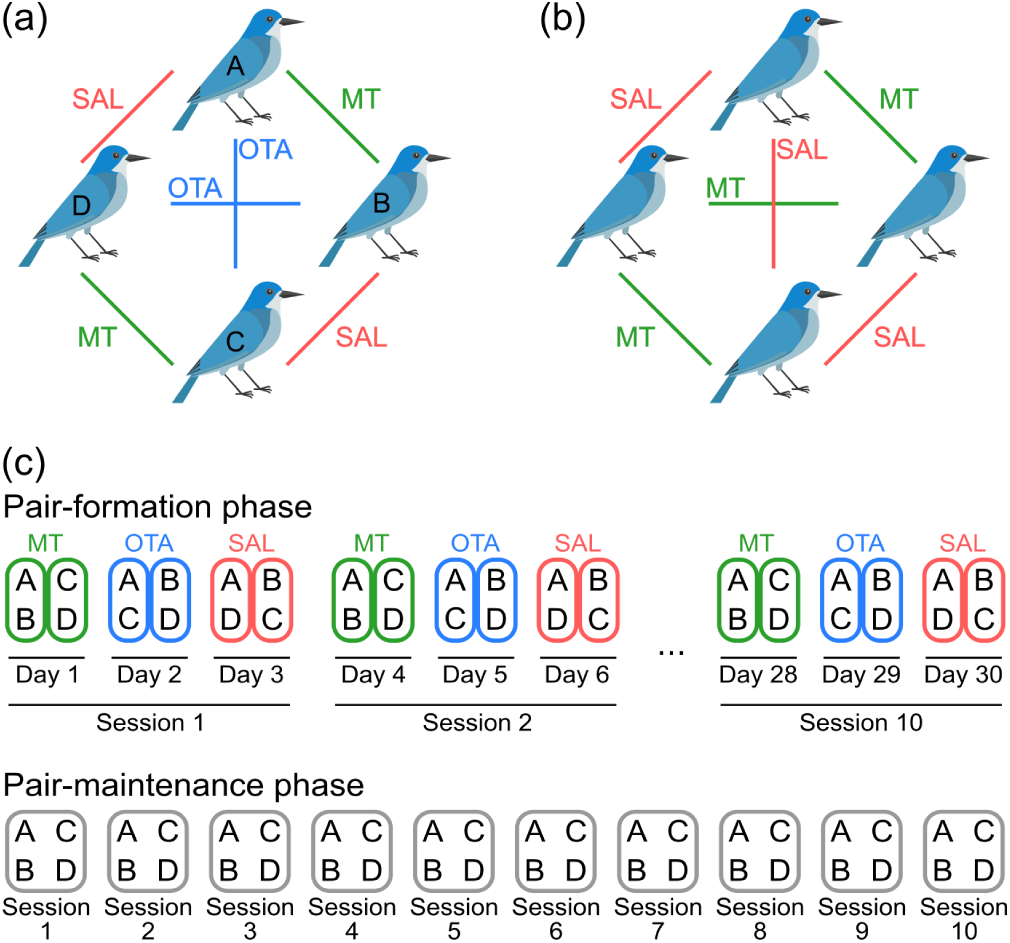
Experimental design. Pinyon jays were repeatedly paired with every other bird from the same squad. We assigned every pair to a hormone condition: (a) Experiment 1 pairs received saline (SAL), mesotocin (MT), or oxytocin antagonist (OTA); (b) Experiment 2 pairs received SAL or MT. (c) Each pair experienced 10 sessions in the pair-formation phase, and each squad experienced 10 sessions in the pairmaintenance phase (A, B, C, and D represent members of a squad). Note: This shows an example of how pairs were run in Experiment 1 for the pair-formation phase with all three treatments. In Experiment 2, pairs were assigned only to SAL or MT.

We assigned each pair of birds a hormone treatment consisting of either saline (SAL), mesotocin (MT), or oxytocin antagonist (OTA; only Experiment 1). Every pair always received the same hormone treatment throughout the duration of the experiment. Because each bird was in three pairs, subjects experienced all hormone conditions, albeit with different partners (Figure 1). In Experiment 1, each individual was assigned one pair for each of the three conditions. In Experiment 2, we simplified the hormonal manipulations by removing the oxytocin antagonist condition, which resulted in each individual involved in either two mesotocin and one saline pairs or one mesotocin and two saline pairs.

### Hormone preparation and administration

We diluted mesotocin (Bachem H2505, Torrance, CA) and oxytocin antagonist (R&D Systems L-368,899, Inc., Minneapolis, MN) to the necessary dose with sterile saline, separated each solution (including the saline control) into individual doses by pipetting 120 microliters into individual microtubes, then froze all samples at -20° C. To ensure experimenters were blind to what hormone corresponded to which condition, we coded all samples as A, B, or C. Doses were calculated per 100 microliters and the additional 20 microliters accounted for any potential spillage. For Experiment 1, the mesotocin dose was 50 micrograms (approximately 24 IU) and oxytocin antagonist was 10 micrograms (based on Smith, Ågmo, Birnie, & French, 2010). Though unclear if related to the mesotocin administration, we observed some unintended side effects during Experiment 1 (e.g., motor-balance irregularities). Further, Duque, Leichner, Ahmann, and Stevens (2018) found a behavioral influence of mesotocin administration using a lower dose, at 30 micrograms per 100 microliters (approximately 14 IU). For these reasons, we reduced the mesotocin dose to 30 micrograms in Experiment 2. To administer a dose, an experimenter used a needle-less syringe to drip the respective solution into the birds’ nostrils. Handling and administration lasted approximately 10-15 seconds per bird.

### Procedure

We sought to manipulate the formation of social bonds by repeatedly pairing birds following exposure to a specific hormone manipulation. Both experiments consisted of three phases: habituation to the testing environment and procedure, a pair-formation phase with repeated sessions of hormone/saline administration for all pairs, and a pair-maintenance phase with repeated sessions of no administration and all four birds of a squad together in a group. Prior to each pair-formation phase session, we administered to each member of a pair its assigned hormone condition (10 sessions for each pair), and all pairs within a squad were cycled through once before repeating any pairs. In summary, there were a total of nine squads (six pairs per squad) comprising of 54 unique pairs and 540 pair-formation sessions overall.

#### Habituation

For habituation sessions, an experimenter transported an individual bird from its home cage to an experimental cage (minimum of 42 × 42 × 60 cm) that had a cup containing the birds’ typical diet. The experimental cage was in another room that was visually isolated from other birds and was the same cage that would later be used during the pair phase. One individual at a time, birds were transported from their home cages, administered a saline dose, then placed in the experimental cage. Each habituation session lasted approximately 15 minutes, and birds were given one session daily for nine weekdays. Thus, all birds were habituated to the testing environment and procedure prior to beginning the pair-formation phase.

#### Pair-formation phase

Pair-formation phase sessions were similar to habituation, except that birds were run in pairs for 45 minutes, and both birds were intranasally administered their preassigned solution immediately prior to being placed in the experimental cage. Specifically, after transporting both birds to the testing room, the experimenter dripped 120 microliters of solution into the birds’ nares, placed both birds in the cage, and immediately exited the room. To distinguish individuals visually, we placed a colored leg band (red, white, blue, or green) on each member of a squad.

Each individual was only run once daily; thus, on any given testing day, only two pairs for each squad were run (i.e., since there are only four birds per squad, running two pairs utilizes all individuals). It took a minimum of three days to cycle through all pairs of each squad. After all pairs were run through a given session, the next block of three sessions began and continued until each bird experienced 10 sessions with each of its three pairs totaling 30 pair-formation sessions per subject. Unlike habituation, we did not introduce food at the beginning of pair phase sessions. However, halfway through Experiment 1 (pair phase sessions 6-10), we introduced a food bowl after 30 minutes to promote interactions between the pair. We discontinued this for Experiment 2 since we observed increased variability in the data following the introduction of food.

#### Pair-maintenance phase

Upon the completion of all pair-formation sessions, we tested each squad in 10 30-minute pair-maintenance phase sessions. In these sessions, we did not administer any solutions, and all four individuals were placed together in a larger cage (66 × 74 × 115 cm). For Experiment 1 only, experimenters introduced two food bowls at the 15-minute mark. We did not introduce any food during Experiment 2 sessions.

### Quantifying pair proximity

We used spatial proximity as our primary measure of social bonding. We also recorded behaviors such as begging, allopreening, food sharing, coordinated displays and calls, aggression, stress panting, and mounting. Because the other behaviors rarely occurred during our sessions, we focus on proximity as our key measure. We video recorded all sessions to precisely measure the distance between the pairs. Coders used Meazure (version 2.0.1, C Thing Software, http://www.cthing.com/Meazure.asp) to capture the coordinates of each bird. Specifically, starting at the 15 s mark and every minute thereafter, we recorded the location of the top-center of each bird’s head, then used those coordinates to calculate the distance between birds for each minute of that session. To account for differences in video size or the camera’s distance from cage, the first recorded point for each session was a fixed, known distance (a horizontal cage bar) which was used to calibrate all following distances for that specific session.

After visualizing and analyzing a subset of Experiment 1 data, we determined that pairs’ mean proximity had stabilized within the first 25 min of each pair session and overall results did not differ between when we analyzed all time points or merely the first 25. Thus, to avoid the increased variability induced by human disturbance and the introduction of food, we only used data from the first 25 min for pair-formation phase sessions. For Experiment 1 pair-maintenance phase sessions, we omitted the proximity data for the minute before, during, and after the experimenter entered the room. Similarly, coders recorded a null measurement whenever the location of a bird’s head was not visible or was unreliable, e.g., when a bird was in mid-flight. All data were scored by one of six coders and, prior to independently coding any sessions, each coder was extensively trained until they reached high reliability. Further, to quantify measurement differences between coders, multiple coders scored the same 48 videos. That is, of the 540 pair-formation sessions, 48 randomly chosen sessions were rated by multiple coders. The remaining 492 sessions were only scored by one of the six coders, with each coder scoring 75-221 sessions.

These data coded by multiple coders were used to calculate the intraclass correlation (ICC) as a measure of inter-rater reliability (Koo & Li, 2016). We calculated ICC from linear mixed models because they directly estimate the variances needed to calculate this measure of repeatability (Nakagawa & Schielzeth, 2010). ICC values greater than 0.9 indicate excellent agreement (Koo & Li, 2016). In an empty, random intercept model, 97.71% [95% CI: 96.36, 98.47] of the variation in pair proximity is accounted for by differences between different videos, suggesting that the different coders shared excellent agreement in quantifying proximities from the same video. We randomly selected one coder’s data for each of the videos for our data analysis.

### Data analysis

We analyzed the data using R (Version 3.6.1; R Core Team, 2019). Data, R code, and supplementary figures and tables are available in the Supplementary Materials and at the Open Science Framework (https://osf.io/67ncp/). The manuscript was created using *rmarkdown* (Version 1.14; Allaire, Xie, McPherson, Luraschi, Ushey, Atkins, Wickham, Cheng, Chang, & Iannone, 2018) and *knitr* (Version 1.23; Xie, 2015), and the reproducible research materials are available from author JRS and at https://osf.io/67ncp/.

#### Model selection

We ran separate analyses of pair proximity for each phase for both experiments (four total datasets), using backward model selection to first find the best-fitting random effect structure, then tested various fixed effects to find the best-fitting model. Different pairs of birds may bond at different rates over time (i.e., across sessions, pairs of birds may move closer to one another as they bond but different pairs may move closer at different rates). For each analysis, we started with the full random effect structure including a random intercept for pair and squad to account for repeated measures, as well as a random slope of pairs over sessions to account for potential differences in how pairs change over time. We sequentially eliminated the weakest, non-significant effects, then ran a nested model comparison (likelihood ratio test) to select the best-fitting random effect structure. A full fixed effect model was then constructed by adding condition (Exp. 1: SAL/MT/OTA; Exp. 2: SAL/MT), session (1-10; centered at final session), their interaction, and the quadratic effect of session. The quadratic fixed effect of session allows for nonlinear change across sessions; i.e., the distance change from one session to next is not consistent. For example, if pairs decrease their distance over the first few sessions but then stay at a fixed distance, this would be a nonlinear relationship that a quadratic term would detect. The final best-fitting model was then selected by sequential deletion and model comparison as detailed above. The significance of terms in all final models was confirmed by Wald tests and non-0 overlapping confidence intervals. Model comparisons are given in Tables S1-S4.

We also calculated Bayes factors (BF) to compare the weight of evidence for alternative models relative to the null (Wagenmakers, 2007). Specifically, we compared each model containing fixed effects to the best-fitting random effect model. We calculated Bayes factors by converting each model’s Bayesian Information Criterion (BIC) using 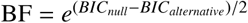 (Wagenmakers, 2007). Bayes factors between 3-10 provide moderate evidence for the alternative hypothesis, those between 10-30 provide strong evidence, those between 30-100 provide very strong evidence, and those above 100 provide extreme evidence (Wagenmakers, Love, Marsman, Jamil, Ly, Verhagen, Selker, Gronau, Dropmann, Boutin, & al., 2018). Reciprocal values (1/3, 1/10, 1/30, 1/100) provide comparable evidence for the null hypothesis.

## Results

### Pair-formation phase

In the pair-formation phase of Experiment 1, we measured the pair proximity for each session and condition. The best-fitting random effect structure included a random intercept for each unique pair and a random slope over sessions; i.e., allowing pairs to change independently over time (random intercept model for pair with versus without random slope: *χ*^2^(2) = 19.98, p < 0.001). However, a random intercept for each squad was not warranted (full versus model without squad: *χ*^2^(1) = 3.26, p = 0.07). Inclusion of condition, session, their interaction, or quadratic effect of session did not improve an empty model (same random effects with no fixed effects, BFs < 0.01). Thus, hormone treatment did not influence pair proximity (Figure 2a).

**Figure 2.**
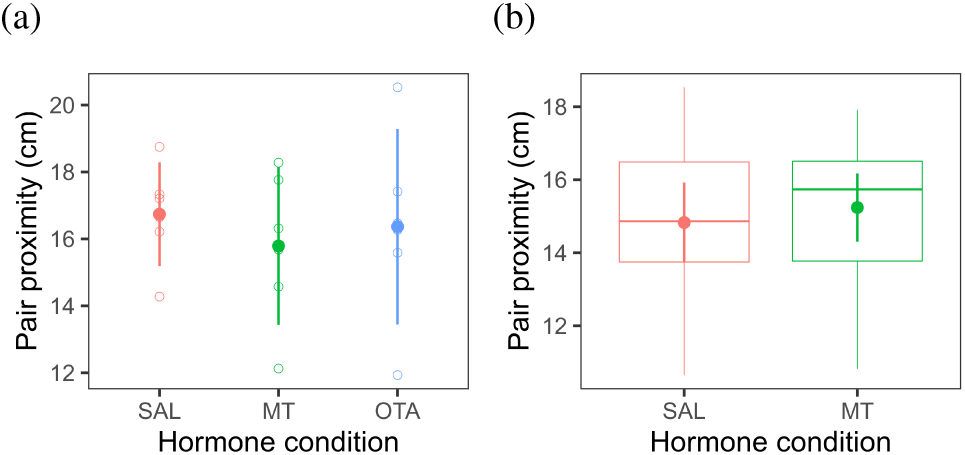
Pair-formation phase pair proximities for each condition for (a) Experiment 1 (6 pairs) and (b) Experiment 2 (12 pairs). Open circles represent individual pairs, horizontal bars represent medians, boxes represent interquartile ranges, whiskers represent full range, closed circles represent means, and error bars represent between-pair confidence intervals. SAL = saline, MT = mesotocin, and OTA = oxytocin antagonist.

In Experiment 2, the best-fitting random effect structure included a random intercept for each unique pair and a random slope over sessions (random intercept model for pair with versus without random slope: *χ*^2^(2) = 22.37, p < 0.001). However, a random intercept for each squad was not warranted (overfit full model versus model without squad: *χ*^2^(1)= 0.00, p > .99). Both linear and quadratic fixed effects of session were warranted (model including linear with versus without quadratic session: *χ*^2^(1) = 7.89, p = 0.005), indicating that pairs perched 0.36 ± 0.12 cm (mean ± standard error) closer each subsequent session, but the decrease in distance diminished by 0.03 ± 0.01 cm each session (Figure S1). That is, though pairs perched more closely over time, the reduction in distance was less pronounced as time progressed. The Bayesian analysis, however, found evidence for no session effect (BF = 0.24). Lastly, inclusion of condition was not warranted (*χ*^2^(1) = 0.35, p = 0.55; Figure 2b).

### Pair-maintenance phase

In the pair-maintenance phase of Experiment 1, the best-fitting random effect structure included only a random intercept for each unique pair (against null model with no random effects; *χ*^2^(1) = 5.05, p = 0.025). A linear fixed effect of session was warranted (against empty model; *χ*^2^(1) = 6.12, p = 0.013), indicating that pairs perched 1.40 ± 0.56 cm closer in each subsequent session (Figure S2). The Bayesian analysis, however, did not find evidence for a session effect (BF = 1.59). No other fixed effects tested (condition or quadratic effect of session) were warranted (Figure 3a).

**Figure 3.**
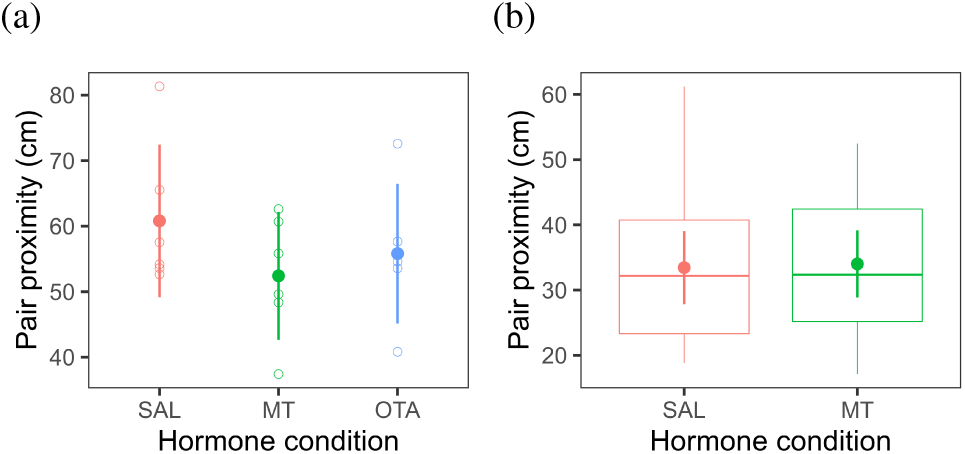
Pair-maintenance phase pair proximities for each condition for (a) Experiment 1 (6 pairs) and (b) Experiment 2 (12 pairs). Open circles represent individual pairs, horizontal bars represent medians, boxes represent interquartile ranges, whiskers represent full range, closed circles represent means, and error bars represent between-pair confidence intervals. SAL = saline, MT = mesotocin, and OTA = oxytocin antagonist.

In Experiment 2, the best-fitting random effect structure included a random intercept for each unique pair and squad but not a random slope (full model with versus without random slope: *χ*^2^(1) = 5.62, p = 0.018). Inclusion of condition, session, their interaction, or quadratic effect of session did not significantly improve an empty model (same random effects with no fixed effects, BFs < 0.09; Figure S2). Thus, hormone condition was not warranted in the best-fitting model (Figure 3b).

As an exploratory analysis of the relationship between pair formation and pair maintenance, we correlated the proximities across the two phases for each pair (Figure S3). Though we do not have evidence for a correlation in Experiment 1 (*r*= -0.06, *p* = 0.81, *BF* = 0.51), we have moderate evidence for a positive correlation between pair formation and pair maintenance phase proximities in Experiment 2 (*r* = 0.42, *p*= 0.01, *BF* = 5.91).

## Discussion

Our analysis of same-sex pinyon jay pairs showed no influence of mesotocin or oxytocin antagonist administration on the proximity of paired birds. Although there was a small effect of session in some models, hormone condition did not influence the proximity of birds for the pair-formation phase or the pair-maintenance phase.

Oxytocin has been implicated in a wide range of social behaviors in mammals (Insel & Young, 2000; Donaldson & Young, 2008), as has isotocin, the oxytocin homologue found in fish (Godwin & Thompson, 2012; Reddon, O’Connor, Marsh-Rollo, Balshine, Gozdowska, & Kulczykowska, 2015) and mesotocin in reptiles (Kabelik & Magruder, 2014). Mesotocin also plays a role in avian maternal care (Chokchaloemwong, Prakobsaeng, Sartsoongnoen, Kosonsiriluk, El Halawani, & Chaiseha, 2013), mating pair bond formation (Pedersen & Tomaszycki, 2012; Klatt & Goodson, 2013), flocking behavior (Goodson, Schrock, Klatt, Kabelik, & Kingsbury, 2009), and prosociality (Duque, Leichner, Ahmann, & Stevens, 2018). Here, we do not demonstrate evidence that mesotocin shapes social bond formation or maintenance in pinyon jays, raising the possibility that mesotocin may function differently than oxytocin. That is, social bonding could be affected differently by mesotocin compared to oxytocin. Though we do not show an effect of mesotocin on bonding in our study, we do not believe that it provides strong evidence against the possibility of mesotocin regulating social bonds in birds for a number of reasons.

Many of the functions of the oxytocin family of peptides are quite evolutionarily conserved, from fish and reptiles to chimpanzees and humans. Though it is possible that functionality may occur in the other species and not birds, this seems unlikely. However, few studies have directly investigated the role of oxytocin-family hormone on social bonds outside of the mating and parenting context. Chimpanzees have higher levels of urinary oxytocin following grooming bouts with socially bonded partners compared to non-bonded grooming partners (Crockford, Wittig, Langergraber, Ziegler, Zuberbühler, & Deschner, 2013; Wittig, Crockford, Deschner, Langergraber, Ziegler, & Zuberbühler, 2014). Yet, this is correlational and only focused on bond maintenance not formation. Administering oxytocin to dogs increases affiliative behaviors to other dogs and humans, but it does not influence spatial proximity and these effects are acute and not long lasting enough to qualify as social bonding (Romero, Nagasawa, Mogi, Hasegawa, & Kikusui, 2014). Female meadow voles do show stronger preferences for familiar partners over unfamiliar partners after oxytocin administration compared to saline, but this effect was measured after only 24 hours (Beery & Zucker, 2010). Though administering oxytocin or mesotocin influences the formation of mating pair bonds (Witt, Carter, & Walton, 1990; Insel & Hulihan, 1995; Pedersen & Tomaszycki, 2012), we do not have strong evidence of these hormones directly shaping formation of same-sex social bonds over time. So it is possible that oxytocin-family hormones facilitate same-sex social bond maintenance but not formation.

It is also possible that mesotocin does facilitate social bond formation, but we simply did not detect it. Though social proximity is generally a good indicator of relationship quality (Croft, Krause, & James, 2008), it may not be a good indicator of the social impact mesotocin has on pinyon jays. It is also possible that behaviors other than proximity are better indicators of social bonds. For pinyon jay mating pairs, proximity is a clear indicator of a pair bond, along with additional behaviors such as begging, allopreening, food sharing, and coordinated displays and calls (Marzluff & Balda, 1992). Unfortunately, we observed very few instances of other behaviors, such as begging, allopreening, aggression, stress panting, and mounting. A more detailed analysis of additional behaviors could reveal differences across hormonal conditions not observed when analyzing social proximity alone.

Our design imposed the pairings of all squads and individuals, rather than allow subjects to have partner choice. Though the freedom to choose partners is clearly important for bonding, similar studies that lack partner choice investigate the relation between oxytocin/mesotocin and social bonds (Williams, Insel, Harbaugh, & Carter, 1994; Beery & Zucker, 2010; Pedersen & Tomaszycki, 2012). In our study, all subjects were adults at the time of capture and thus likely experienced normal social interactions up to that point. In captivity, physical contact has been limited in our birds, but they have been surrounded by other individually housed birds, and most pairs in our study were from the same housing room. Therefore, most pairs should have been familiar with each other prior to the study. Nevertheless, the lack of direct physical contact with others during captivity could have affected how these birds respond to mesotocin administration and form new bonds. Most studies on social history, housing style, and their effects on adult behavior focus on a few key rodent and primate species, and overall findings suggest these effects are most pronounced when the isolation occurs early in development (Lickliter, Dyer, & McBride, 1993; Carere, Welink, Drent, Koolhaas, & Groothuis, 2001; Olsson & Westlund, 2007). Among adults, social instability (e.g., removal from a pre-existing group) is broadly stressful and can impact how individuals interact in the group (Smith, Birnie, & French, 2011; Stocker, Munteanu, Stöwe, Schwab, Palme, & Bugnyar, 2016; Munteanu, Stocker, Stöwe, Massen, & Bugnyar, 2017). Despite these observations, differences in captive housing style do not always result in demonstrable changes in behavior, as compared to wild conspecifics (Gazes, Brown, Basile, & Hampton, 2013). Additionally, we have used these wild caught, individually housed adult pinyon jays to study various social behaviors, including dominance interactions involving competition over food (Bond, Kamil, & Balda, 2003; Paz-y-Miño C, Bond, Kamil, & Balda, 2004), food sharing between jays that had never been paired (Duque & Stevens, 2016), and mesotocin effects on prosocial decision making (Duque, Leichner, Ahmann, & Stevens, 2018). Finally, our birds were at least 6 years old when tested. Though many same-sex social bonds may form earlier than this in the wild, the annual influx of new birds into pinyon jay flocks combined with the flexible social structure that they experience suggests that these bonds may also form later in life. Nevertheless, the social environment likely moderates the effect of hormones on social bonds.

Additionally, insufficient dosage or sub-optimal timing of the dosage may have interfered with the establishment of the social bonds. We used dosages based on our previous study showing acute effects of mesotocin on prosocial food sharing (Duque, Leichner, Ahmann, & Stevens, 2018). However, it is possible that different dosages are required to induce the longer-term effects on social bonds. It is also possible that the immediate time course of administration and behavioral testing did not match that needed to establish the bonds. In our design, birds received one hormone dose and were placed together in a cage for 45 minutes. For a given pair, this occurred roughly every three days and each individual experienced ten sessions with each of its partners. The duration and frequency of social interactions experienced in the lab likely differ from that experienced in the wild (Marzluff & Balda, 1992). Nonetheless, in prior studies in which partner pairs have been imposed and not chosen, these jays preferentially shared food with specific individuals and rarely with others (Duque & Stevens, 2016) and increased the proportion of choices a jay will make to feed a partner over an empty cage when administered mesotocin (Duque, Leichner, Ahmann, & Stevens, 2018). Finally, each pair experienced ten sessions with each partner. Some of the statistical models showed effects of sessions on proximity, with pairs getting closer over time. Though they did not differ across hormone treatment, it is possible that we did not give the bonds enough time to form, and additional treatments and sessions are needed to build the bonds. Thus, it is possible our mesotocin impacts birds in ways that were not captured by our specific measures or study design.

While we chose to investigate the effects of mesotocin, it is plausible that other hormones may play a stronger role in avian social bonding. For instance, both administration of vasotocin (the avian homologue of the mammalian arginine vasopressin) as well as neural vasotocin activity is related to gregariousness in zebra finch, but the effect is most evident in males (Goodson, Lindberg, & Johnson, 2004; Goodson, Schrock, Klatt, Kabelik, & Kingsbury, 2009). Importantly, vasotocin promoted a preference for a larger flock size in male zebra finch, but did not impact the amount of time spent in close proximity (Kelly, Kingsbury, Hoffbuhr, Schrock, Waxman, Kabelik, Thompson, & Goodson, 2011). Thus, the role of vasotocin in pinyon jay social behavior warrants investigation. Further, low sample size prevents our testing of sex differences, but it is possible that mesotocin or vasotocin impacts the sexes differently.

Lastly, the level of circulating hormones is only one way in which hormones might regulate social bond formation. It is unclear how measurements and administration of oxytocin-family hormones outside of the brain relate to levels in the brain (McCullough, Churchland, & Mendez, 2013; Evans, Dal Monte, Noble, & Averbeck, 2014), particularly in corvids, among which relatively little mesotocin research has been conducted (Duque, Leichner, Ahmann, & Stevens, 2018). Nevertheless, there is evidence in other species of peripheral levels correlating with social behavior (Crockford, Wittig, Langergraber, Ziegler, Zuberbühler, & Deschner, 2013; Wittig, Crockford, Deschner, Langergraber, Ziegler, & Zuberbühler, 2014) and peripheral administration influencing social behavior (Smith, Ågmo, Birnie, & French, 2010; Romero, Nagasawa, Mogi, Hasegawa, & Kikusui, 2014). Yet, individuals also vary in their underlying sensitivity to those hormones, primarily determined by the number and distribution of the receptors to which those hormones bind. For example, differences in the density of oxytocin/vasopresson neurons in the brain underlie whether a prairie vole will form a monogamous bond with its partner, or be polygamous (Insel, Winslow, Wang, & Young, 1998). Thus, it would be highly informative to analyze the localization of mesotocin receptors across the pinyon jay brain to shed light on what makes this particular species remarkably social, as compared to even its closest sister species (Marzluff & Balda, 1992).

Though hormone administration did not influence pair formation or maintenance in this study, we did find evidence that pair formation in the first phase correlated with pair maintenance in the second phase in Experiment 2 (Figure S3). This finding suggests that being paired in dyads did in fact create social bonds that carried over into a larger social network. Thus, we have validated an experimental paradigm to explore pair formation and maintenance. However, this effect may be magnified by the housing environment (individually housed birds) and may differ from situations in which subjects can voluntarily choose social partners. We did not find evidence for a correlation in Experiment 1, but the smaller sample size precludes us from supporting the presence or absence of a correlation.

Here, we find that administration of mesotocin or oxytocin antagonist did not impact how closely two previously unfamiliar birds perched next to one another. However, future investigations are warranted to clarify whether mesotocin influences (1) other forms of behaviors during bond formation and the time course of those effects, (2) the relationship between administered mesotocin and circulating levels in the brain, (3) the role of related hormones (e.g., vasotocin), and (4) the role of mesotocin on social behaviors in other corvid species. Given the variation in levels of sociality and cooperation across corvids, exploring the hormonal and neural underpinning of these behaviors could provide valuable insights into the evolution and mechanisms of social behavior.

## Acknowledgments

This research was supported, in part, by a Nebraska EPSCoR FIRST Award and a University of Nebraska-Lincoln Layman Award to J.R.S. and a National Science Foundation Graduate Research Fellowship Program award (DGE-10410000) to J.F.D.

We would like to thank the undergraduate research assistants Megan Bosworth, Allie Cruikshank, Gage Grutz, Marisa Howell, Gretchen Lusso, Maddie Mathias, and Elise Thayer for collecting the data, laboratory technician Jesse Baumann for maintaining the bird colony, and Jeffrey French and Aaryn Mustoe for advice on mesotocin administration.

