## Supplementary Figures and Tables for "The role of mesotocin on social bonding in pinyon jays"

**Citation:** Duque, J.F., Rasmussen, T., Rodriguez, A., & Stevens, J.R. (in press). The role of mesotocin on social bonding in pinyon jays. *Ethology*.

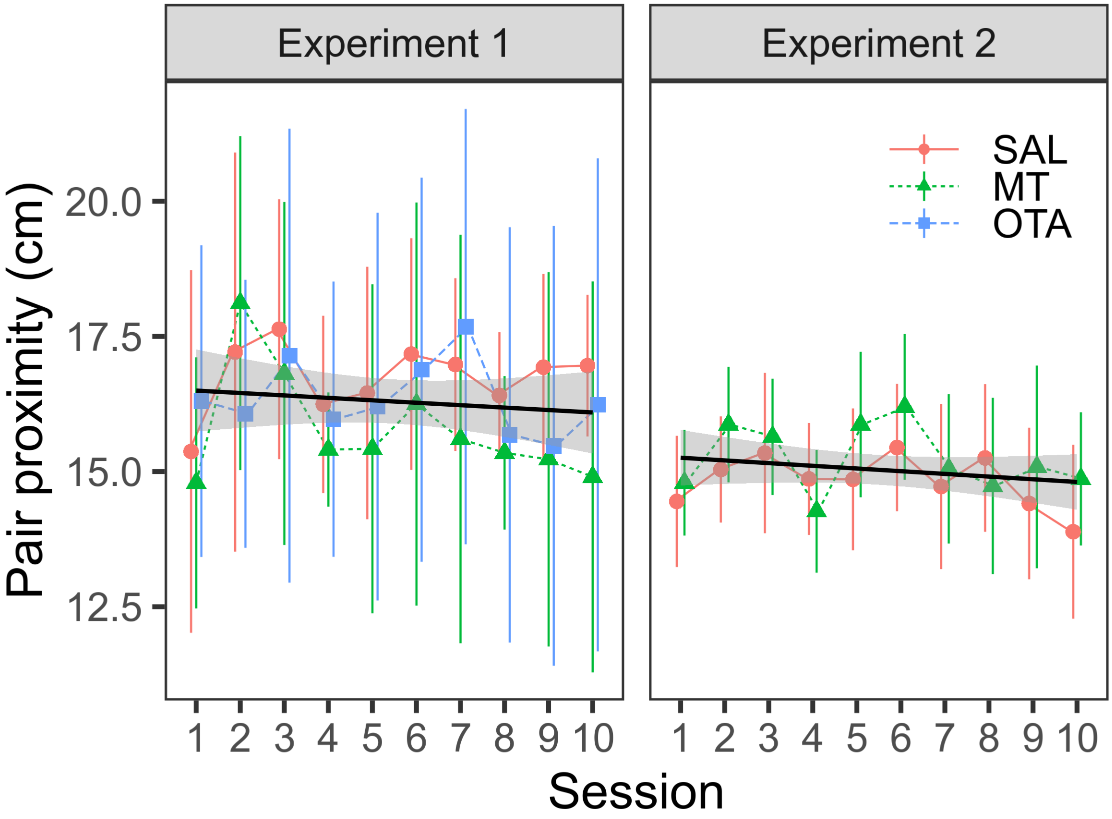


*Figure S1*. Pair-formation phase pair proximities for each session and condition in Experiment 1 (18 pairs) and Experiment 2 (36 pairs). Experiment 1 showed no effect of hormone condition or session on distance, but Experiment 2 showed a decrease in distance over session. Symbols represent means, error bars represent between-pair confidence intervals.

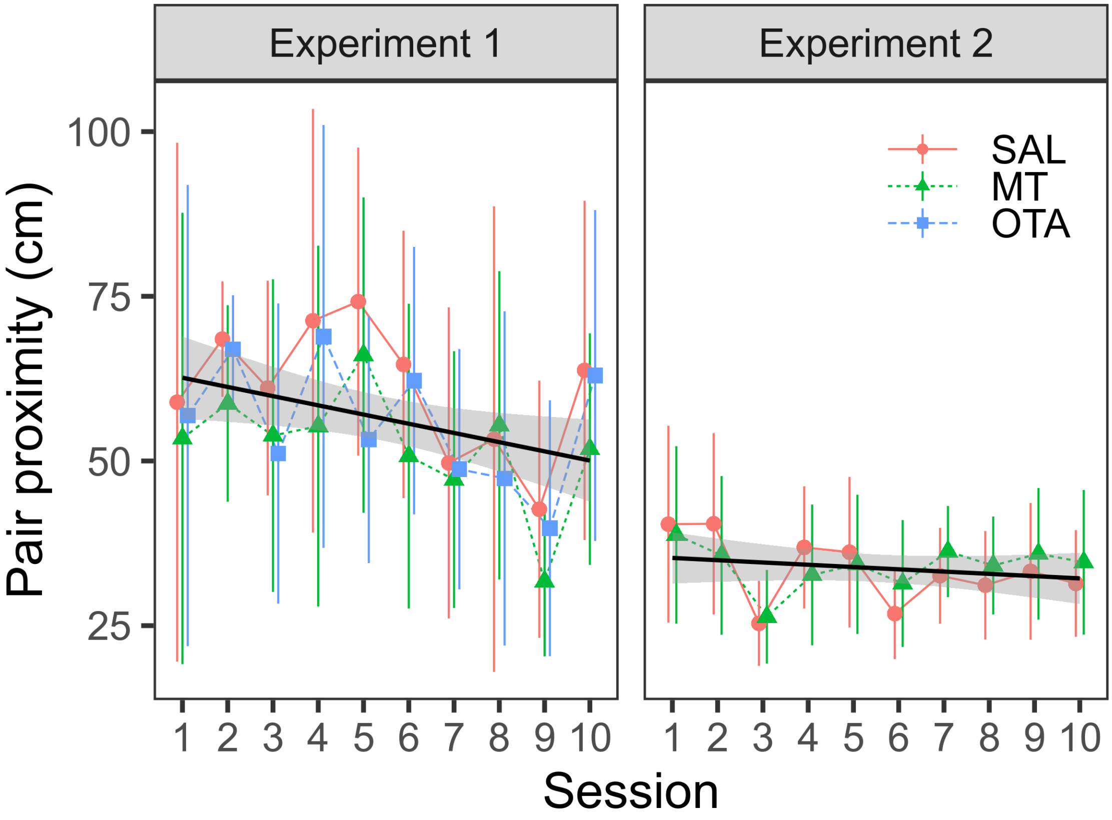


*Figure S2.* Pair-maintenance phase pair proximities for each session and condition in Experiment 1 (18 pairs) and Experiment 2 (36 pairs). In Experiment 1, pair distances decreased over session, but hormone condition did not influence distances. In Experiment 2, neither hormone condition nor session influenced distance. Symbols represent means, error bars represent between-pair confidence intervals.

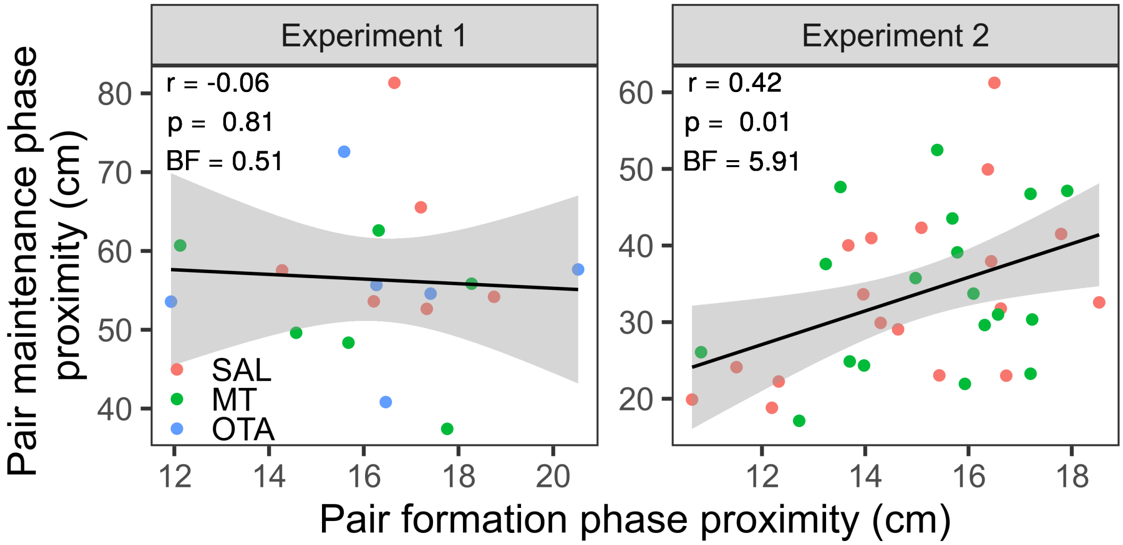


*Figure S3*. Correlations of pair proxmimities across pair-formation and pair-maintenance phases in Experiment 1 (18 pairs) and Experiment 2 (36 pairs). In Experiment 1, we do not have evidence that pair proximities correlate across phases, but, in Experiment 2, proximities positively correlated across phases.

Table S1

*Experiment 1 Pair-Formation Phase Model Comparison*

| | | | Model fit | | | | Likelihood Ratio Tests | | | |
|---|---|---|---|---|---|---|---|---|---|---|
| Model specification | Random effects | Fixed effects | AIC | BIC | logLik | df | $\chi^2$ | df | p-value | BF |
| Random effects models | | | | | | | | | | |
| Pair | (1 \| pair) | - | 797.19 | 806.77 | -395.60 | 3 | | | | |
| Pair + session slope | (1 + session0 \| pair) | - | 781.21 | 797.18 | -385.61 | 5 | 19.98 | 2 | 0.000 | |
| Group + pair + session slope | (1 \| group) + (1 + session0 \| pair) | - | 779.96 | 799.12 | -383.98 | 6 | 3.26 | 1 | 0.071 | |
| Fixed effects models | | | | | | | | | | |
| RE only | (1 + session0 \| pair) | - | 781.21 | 797.18 | -385.61 | 5 | | | | |
| Session | (1 + session0 \| pair) | session0 | 782.93 | 802.08 | -385.46 | 6 | 0.29 | 1 | 0.592 | 0.007 |
| Condition | (1 + session0 \| pair) | condition | 784.85 | 807.20 | -385.43 | 7 | 0.07 | 1 | 0.787 | 0.007 |
| Condition + session | (1 + session0 \| pair) | condition + session0 | 786.57 | 812.11 | -385.28 | 8 | 0.29 | 1 | 0.592 | 0.001 |
| Condition + session + session squared | (1 + session0 \| pair) | condition + session0 + session0^2 | 786.26 | 814.99 | -384.13 | 9 | 2.31 | 1 | 0.129 | 0.000 |
| Condition * session + session squared | (1 + session0 \| pair) | condition * session0 + session0^2 | 789.20 | 824.32 | -383.60 | 11 | 1.06 | 2 | 0.588 | 0.000 |

Table S2

*Experiment 2 Pair-Formation Phase Model Comparison*

| | | | Model fit | | | | Likelihood Ratio Tests | | | |
|---|---|---|---|---|---|---|---|---|---|---|
| Model specification | Random effects | Fixed effects | AIC | BIC | logLik | df | $\chi^2$ | df | p-value | BF |
| Random effects models | | | | | | | | | | |
| Pair | (1 \| pair) | - | 1557.71 | 1569.37 | -775.85 | 3 | | | | |
| Pair + session slope | (1 + session0 \| pair) | - | 1539.34 | 1558.77 | -764.67 | 5 | 22.37 | 2 | 0.000 | |
| Group + pair + session slope | (1 \| group) + (1 + session0 \| pair) | - | 1541.34 | 1564.66 | -764.67 | 6 | 0.00 | 1 | 1.000 | |
| Fixed effects models | | | | | | | | | | |
| RE only | (1 + session0 \| pair) | - | 1539.34 | 1558.77 | -764.67 | 5 | | | | |
| Session | (1 + session0 \| pair) | session0 | 1540.28 | 1563.59 | -764.14 | 6 | 1.07 | 1 | 0.301 | 0.090 |
| Session + session squared | (1 + session0 \| pair) | session0 + session0^2 | 1534.39 | 1561.59 | -760.19 | 7 | 7.89 | 1 | 0.005 | 0.245 |
| Condition + session + session squared | (1 + session0 \| pair) | condition + session0 + session0^2 | 1536.04 | 1567.12 | -760.02 | 8 | 0.35 | 1 | 0.553 | 0.015 |
| Condition * session + session squared | (1 + session0 \| pair) | condition * session0 + session0^2 | 1538.00 | 1572.97 | -760.00 | 9 | 0.04 | 1 | 0.848 | 0.000 |

Table S3

*Experiment 1 Pair-Maintenance Phase Model Comparison*

| | | | Model fit | | | | Likelihood Ratio Tests | | | |
|---|---|---|---|---|---|---|---|---|---|---|
| Model specification | Random effects | Fixed effects | AIC | BIC | logLik | df | $\chi^2$ | df | p-value | BF |
| Random effects models | | | | | | | | | | |
| Empty | 1 | - | 1644.98 | 1651.36 | -820.49 | 2 | | | | |
| Pair | (1 \| pair) | - | 1641.93 | 1651.51 | -817.97 | 3 | 5.05 | 1 | 0.025 | |
| Group + pair | (1 \| group) + (1 \| pair) | - | 1643.15 | 1655.93 | -817.58 | 4 | 0.78 | 1 | 0.377 | |
| Group + pair + session slope | (1 \| group) + (1 + session0 \| pair) | - | 1646.52 | 1665.68 | -817.26 | 6 | 0.63 | 2 | 0.728 | |
| Fixed effects models | | | | | | | | | | |
| RE only | (1 \| pair) | - | 1641.93 | 1651.51 | -817.97 | 3 | | | | |
| Session | (1 \| pair) | session0 | 1637.81 | 1650.58 | -814.91 | 4 | 6.12 | 1 | 0.013 | 1.592 |
| Condition + session | (1 \| pair) | condition + session0 | 1639.51 | 1658.67 | -813.76 | 6 | 2.30 | 2 | 0.317 | 0.028 |
| Condition * session | (1 \| pair) | condition * session0 | 1643.47 | 1669.01 | -813.73 | 8 | 0.04 | 2 | 0.980 | 0.000 |
| Condition * session + session squared | (1 \| pair) | condition * session0 + session0^2 | 1644.95 | 1673.68 | -813.47 | 9 | 0.52 | 1 | 0.469 | 0.000 |

Table S4

*Experiment 2 Pair-Maintenance Phase Model Comparison*

| | | | Model fit | | | | Likelihood Ratio Tests | | | |
|---|---|---|---|---|---|---|---|---|---|---|
| Model specification | Random effects | Fixed effects | AIC | BIC | logLik | df | $\chi^2$ | df | p-value | BF |
| Random effects models | | | | | | | | | | |
| Empty | 1 | - | 3187.05 | 3194.82 | -1591.53 | 2 | | | | |
| Pair | (1 \| pair) | - | 3155.75 | 3167.41 | -1574.87 | 3 | 33.30 | 1 | 0.000 | |
| Group | (1 \| group) | - | 3149.62 | 3161.28 | -1571.81 | 3 | 6.13 | 0 | 0.000 | |
| Group + pair | (1 \| group) + (1 \| pair) | - | 3146.00 | 3161.55 | -1569.00 | 4 | 5.62 | 1 | 0.018 | |
| Group + pair + session slope | (1 \| group) + (1 + session0 \| pair) | - | 3147.89 | 3171.21 | -1567.95 | 6 | 2.11 | 2 | 0.348 | |
| Fixed effects models | | | | | | | | | | |
| RE only | (1 \| group) + (1 \| pair) | - | 3146.00 | 3161.55 | -1569.00 | 4 | | | | |
| Session | (1 \| group) + (1 \| pair) | session0 | 3146.92 | 3166.35 | -1568.46 | 5 | 1.08 | 1 | 0.298 | 0.091 |
| Session + session squared | (1 \| group) + (1 \| pair) | session0 + session0^2 | 3146.14 | 3169.45 | -1567.07 | 6 | 2.78 | 1 | 0.095 | 0.019 |
| Condition + session + session squared | (1 \| group) + (1 \| pair) | condition + session0 + session0^2 | 3148.09 | 3175.29 | -1567.04 | 7 | 0.05 | 1 | 0.824 | 0.001 |
| Condition * session + session squared | (1 \| group) + (1 \| pair) | condition * session0 + session0^2 | 3148.53 | 3179.62 | -1566.26 | 8 | 1.56 | 1 | 0.212 | 0.000 |
